# Estimating probabilistic dark diversity based on the hypergeometric distribution

**DOI:** 10.1101/636753

**Authors:** Carlos P. Carmona, Robert Szava-Kovats, Meelis Pärtel

## Abstract

1. The biodiversity of a site includes the absent species from the region that are theoretically able to live in the site’s particular ecological conditions. These species constitute the dark diversity of the site. Unlike present species, dark diversity is unobservable and can only be estimated. Most existing methods to designate dark diversity act in a binary fashion. However, dark diversity is more suitably defined as a fuzzy set—in which the degree of certainty about species membership is expressed as a probability.
2. We present a new method to estimate probabilistic dark diversity based on the hypergeometric distribution. The method relies on co-occurrences to infer the strength of the association between pairs of species and assign probabilistic adscription to dark diversity to absent species. We compare it with two established methods to estimate dark diversity (Beals index and favorability correction). To test the methods, we created simulations based on individual agents in which the suitability of each species in each site is known. We compared the ability of the methods to accurately predict suitability and the size of dark diversity, and compared their sensitivity to data availability. Further, we assessed the methods in two real datasets with nested sampling designs.
3. Our simulations revealed that predictions of the Beals method were extremely sensitive to species frequency, and predicted suitability poorly. The Favorability transformation corrected this relationship, but did still predicted extremely low probabilities for species with very little information. The Hypergeometric method outperformed the Beals and Favorability methods in all considered aspects in the simulations and displayed better characteristics in the real datasets.
4. Probabilistic consideratiosn of biodiversity will help to acknowledge the uncertainty associated with ecological information. Although the Beals method has been described as the best estimator of dark diversity, it should be preferred only when the goal is to predict future apperances of species. However, studies on dark diversity should focus on the ecological affinities of species. The Hypergeometric method is the most promising method to estimate probabilistic dark diversity and species pool composition based on co-occurrences.

## Introduction

The biodiversity of a site consists not only of those species actually present, but also of absent species from the region that are theoretically able to live in the site’s particular ecological conditions (its dark diversity; Pärtel, Szava-Kovats, & Zobel, 2011). Unlike present species, dark diversity is, by definition, unobservable and must be estimated. The increasing recognition of the importance of considering absent species (Bennett & Pärtel, 2017; de Bello et al., 2012; Pärtel et al., 2011) has recently seen the development of methods to estimate the size and composition of dark diversity (de Bello et al., 2016; Karger et al., 2016; Lewis, Szava-Kovats, & Pärtel, 2016), although ample room remains for methodological improvements. Methods to estimate dark diversity include the use of indicators of the position of species’ niches along environmental gradients (de Bello et al., 2016; Lewis et al., 2017), species distribution modelling (Estrada, Barbosa, & Real, 2018; Ronk, de Bello, Fibich, & Pärtel, 2016), regional surveys of the habitat of interest (Jiménez-Alfaro et al., 2018), or species co-occurrence patterns (Brown et al., 2019; de Bello et al., 2016; Lewis et al., 2016).

Many of these methods designate dark diversity in a binary fashion, i.e., any given species either belongs (1) or does not belong (0) to local dark diversity. However, binary classification requires establishing thresholds to define which species are included in dark diversity. Despite efforts to make this procedure as aseptic as possible, the selection of thresholds remains rather arbitrary (Karger et al., 2016), can affect the results (Lewis et al., 2016), and is often difficult to justify. By contrast, dark diversity is more suitably defined as a fuzzy set—in which the degree of certainty about species membership is expressed as a probability—rather than as a binary designation. In other words, the probability of a species passing through all the different ecological filters ultimately determines the probability that the species is part of the dark diversity of a given site.

Although the justification for probabilistic approaches to dark diversity is long recognized (e.g. Mokany & Paini, 2011; Pärtel, Zobel, Zobel, van der Maarel, & Partel, 1996), methods adopting this approach are only recently being developed (Karger et al., 2016; Lessard et al., 2016; Real, Márcia Barbosa, & Bull, 2017). Species co-occurrence patterns offer a pragmatic method for the probabilistic approach. Species that frequently co-occur share similar ecological requirements (integrating both abiotic and biotic conditions). Imagine we are interested in the status of a particular species that has not been observed in a community. The presence of other species that tend to be found together with this species suggests that the probability of membership in local dark diversity is high. The most widely used method to estimate dark diversity based on co-occurrence patterns is the Beals index (Beals, 1984; Ewald, 2002). Evidence suggests that estimations of dark diversity based on the Beals index have greater predictive ability than relying on databases with habitat requirements of species (de Bello et al., 2016; Lewis et al., 2016). This method assigns to each species and site the probability of the species being present, which is computed by combining information on the identity of the species actually found in the community (observed diversity) and their patterns of co-occurrence with the focal species. However, Beals values increase monotonically with the frequency of the species in the region (De Cáceres & Legendre, 2008; Lewis et al., 2016; Münzbergová & Herben, 2004). This is problematic, because the fact that a species is rarely observed in a set of communities is not necessarily an indicator that the species is not part of the dark diversity of some sites, particularly if dispersal limitation plays a role (Jiménez-Alfaro et al., 2018; Riibak et al., 2015). Actually, the probability that a species will appear in a site where it is currently absent depends on a combination of the suitability of the local conditions and factors related to dispersal, including regional frequency and dispersal ability. Accordingly, Beals values have been used in studies aiming to predict species appearances in the near future without distinguishing habitat suitability per se (Karger et al., 2016). However, when studying dark diversity we are interested only on species suitability. One way to resolve this issue is to apply species-specific thresholds (Münzbergová & Herben, 2004), resulting in a binary classification of species. Although such a classification is independent of species frequency, it lacks the preferred notion of dark diversity in probabilistic terms.

One alternative is to transform indices affected by species frequency (such as Beals) into pure indicators of the suitability of the local conditions for each particular species (Favorability; Real, Barbosa, & Vargas, 2006). The favorability transformation provides information on the likelihood of a species to be found in a site with respect to random expectations (i.e. regardless of its presence/absence ratio in the dataset; Real et al., 2006). This solution—which has been applied to logistic regressions in the context of species distribution modelling (Olivero et al., 2017; Real et al., 2017)—could also be applied to estimate probabilistic dark diversity from the Beals index. Alternatively, rather than solving the issue of frequency with post-hoc transformations, we propose that species suitability in a site can be estimated directly by comparing the realised co-occurrence patterns of each pair of species to that expected under the assumption of their complete lack of association. The degree to which the observed co-occurrence between a pair of species departs from random association can be then used as the indicator value for that pair of species. Associations between pairs of species can be analysed using the hypergeometric distribution (Griffith, Veech, & Marsh, 2016).

In this paper, we advance towards the establishment of methods to estimate probabilistic dark diversity using species co-occurrence matrices. We first present a novel method based on the hypergeometric probability distribution to assign probabilistic estimates of dark diversity. We test this, along with raw Beals values and its transformation into favorability, in a simulated dataset created through individual-based modelling, resulting into communities with known observed and dark diversity. We subsequently compare the different methods using a real dataset with a nested sampling structure (Lewis et al., 2016). This comparison allows us to distinguish features of these methods, including their probabilistic distributions, their ability to estimate accurately the ecological suitability of sites for species, or their dependence on the amount of data available.

### Simulations and dark diversity estimations

Comparing the performance of methods to estimate dark diversity is challenging because dark diversity is not observable in natural conditions. Some studies have used datasets with nested hierarchical sampling designs, where vegetation is sampled in a small plot that is contained within a larger plot (Brown et al., 2019; de Bello et al., 2016; Lewis et al., 2016). In these studies, the information from the smaller plot is used to build a species x species co-occurrence matrix, and the estimations of dark diversity made from the smaller plots are confronted with the species present in the larger plots. It is unclear, however, to what point species in the larger plot reflect the true dark diversity of the smaller plot. Species whose ecological requirements match those of the site are not necessarily present in the surroundings. This can happen, for example, when a species has been unable to disperse to a favourable site, which is more likely the case for regionally rare species. As a result, considering that the dark diversity of the small plot can be derived from the species present in the surroundings likely favours methods whose predictions reflect species frequency. However, as discussed above, these methods do not necessarily reflect better the suitability of species. Simulations which assign the match between the ecological requirements of species and the environmental characteristics of sites are a valuable alternative in this case (Lewis et al., 2016). In short, we created a virtual landscape containing different habitats and a set of species with different suitability for these habitats and allowed communities to develop following simple rules for a period of time (see below). Finally, we sampled the communities and used the co-occurrence pattern of species to estimate dark diversity with the different methods, which we finally compared with species suitability.

Simulations were based on Jõks & Pärtel (2019), with the difference that our agents represented individuals of a species rather than populations. We created a 100 x 100 grid divided into 100 plots (each encompassing 10 x 10 cells); cells could either contain an individual or be empty. Individuals acted according to simple rules that corresponded to some of the basic processes that determine diversity (selection, drift, and dispersal; see below and Vellend, 2010). Among these processes, selection depended on the suitability of each species to each plot. For this, we assigned the same value for environment to all the cells in the same plot, which was drawn from a normal distribution with µ=0 and σ=5. We then created a set of 100 species, with each species having an optimal value in the environment drawn from a uniform distribution from -10 to 10; all individuals of a species had the same value (i.e. there was no intraspecific variability). Once these values were assigned, we estimated the distance between each community’s environment and each species optimum, considering the environment as a circular variable. Suitability indicates how close an environment is to the optimum of a given species; suitability was 1 when the environment value in the plot was equal to the species optimum and decreased towards 0 as distance increased (following a normal distribution).

Simulations started with an empty grid (no individuals present), and were run for 5250 sequential cycles. In each cycle, the following processes (and sub-processes) took place:

#### Dispersal

Species were added to communities through dispersal (Vellend, 2010), which had two sources in our simulation: immigration from the region and reproduction. Immigration simulated the arrival to the grid of individuals belonging to species from outside the landscape. In each cycle, each cell had a 10% probability of receiving an individual from a randomly selected species from the region. Established individuals (see “Selection” below) had a 40% probability of reproducing; reproducing individuals created a propagule which was dispersed in a random direction at a distance that was chosen from a log-normal distribution with a mean value of 10% the maximum distance between cells in the grid. All species had similar dispersal abilities. To avoid edge effects, we set periodic boundary conditions in the grid; this way, when a propagule reached the boundaries of the grid, its dispersal continued from the opposite side. When individuals from more than one species arrived at the same cell in a cycle, the retained species was randomly selected among the arriving species.

#### Selection

This category included processes regulating interactions between species and of species with their environment. We considered two main selection sub-processes, both related with suitability: establishment and competition. Establishment decided whether a propagule arriving to a cell formed an adult individual or died. The probability that a propagule established in a cell was equal to the suitability of the species in the corresponding plot. Competition took place when an individual was able to establish in a cell previously occupied by another individual (the “local” individual). In this case, the difference in competitive abilities between the arriving and the local individuals was estimated as their difference in suitability (Diff_Suit_ = suitability_local_ – suitability_dispersed_). The probability that the local would persist was estimated as the logistic function of Diff_Suit_. Through the combined effect of establishment and competition, species with higher suitability for a given plot should be more frequent and abundant in this plot.

#### Drift

This category included processes that randomly changed species abundances (Vellend, 2010). We incorporated it in the simulations by including mortality: in each cycle, each individual had a fixed 10% probability of dying, regardless of its suitability.

We built a species x species co-occurrence matrix from the composition after the final cycle, and then estimated probabilistic dark diversity for each plot using three different co-occurrence based methods: the Beals index, Favorability, and the newly developed Hypergeometric method. We provide the functions for each of these processes, as well as the code used for the simulations in Appendix 1.

### Probabilistic estimations of dark diversity

#### HYPERGEOMETRIC METHOD

The premise of the hypergeometric method is simple: for each pair of species we can compare their realised number of co-occurrences with random expectations (i.e. if there was no association between species). Let us consider two species *i* and *j*; the probability that they co-occur in a number of sites *M* is given by the mass function of the hypergeometric distribution (Griffith et al., 2016; Veech, 2013):

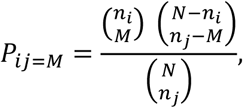

where *n*_*i*_ and *n*_*j*_ are the total number of occurrences of species *i* and *j*, respectively, and *N* is the total number of sites sampled. The mean of this distribution 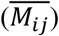 denotes the expected number of co-occurrences between species *i* and *j* is given by:

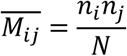

Logically, if the number of actual co-occurrences is greater than expected by chance, the two species are positively associated, and vice versa. We can estimate this departure from expected (ES, effect size) simply by subtracting 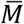 to *M*:

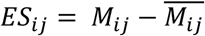

*ES*, however does not convey information on the strength of the association (or lack thereof) between two species. For this, we can estimate standardized effect sizes (SES) by dividing the effect size by the square root of the variance of the hypergeometric distribution (the standard deviation):

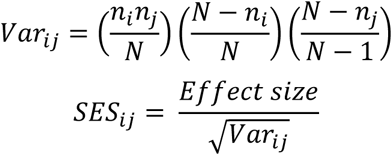

SES indicates how many standard deviations the observed number of co-occurrences is from the expected value. They can then be expressed as probabilities (*P*_*ij*_) by confronting the SES value with the cumulative normal distribution function with mean=0 and standard deviation=1. Probabilities close to 1 indicate that the two species are positively associated, whereas probabilities close to 0 indicate that the two species are negatively associated; intermediate values denote a random association. This procedure can be applied to all pairs of species to build a symmetric indication matrix reflecting the strength of the association between all species pairs. The indication matrix can then be used to predict the probabilistic dark diversity of a given site (*k*) for which we know the observed diversity. This probability can be estimated for each of the absent species in the site (i.e. all species in the dataset that were not present in the site) simply by averaging the indication values of the species actually present in the community:

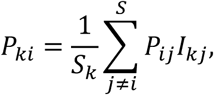

where *S*_*k*_ is the total number of species found in site *k, I*_*kj*_ reflects the incidence (0, 1) of the indicator species *j* in site *k*, and *S* is the total number of species in the region. Hence, the probability of an absent species belonging to the dark diversity of a site is high if it tends to have positive associations with those species that are present, and negative associations result in a low probability of membership.

#### BEALS INDEX

The Beals probability that a species *i* should be present in a site *k* (*P*_*ki*_) can be estimated following (Münzbergová & Herben, 2004):

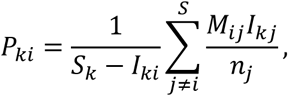

where *S_k_* is the total number of species found in site *k, I*_*ki*_ and *I*_*kj*_ reflect the incidence (0, 1) of species *i* and *j* in site k, respectively, *S* is the total number of species in the region, *M*_*ij*_ is the number of co-occurrences between species *i* and *j*, and *n*_*j*_ is the total number of occurrences of species *j*, considering all sites. The probabilities predicted by the Beals index are correlated with the frequency of the species in the considered dataset, which has led some authors to recommend setting a species-specific probability threshold, which effectively creates a binary index (Lewis et al., 2016; Münzbergová & Herben, 2004).

#### FAVORABILITY INDEX

An alternative that avoids thresholding and makes the probabilities independent of species frequency is the favorability index proposed by Real, Barbosa, & Vargas (2006):

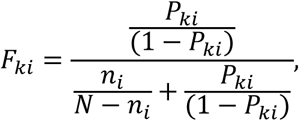

where *F*_*ki*_ is the favorability of site *k* for species *i, P*_*ki*_ is a probability index affected by the global frequency of the species (i.e. the Beals index in this case).

### Methods performance comparison

The advantage of using simulations to test dark diversity methods is that information about the suitability of absent species in each plot is predetermined and can be compared to the probabilities obtained from each method. We designed different tests to compare specific aspects of the methods.

#### TEST 1: CORRELATION WITH SUITABILITY AND BIAS

##### Test rationale

We predicted the probabilities of all the absent species from all the communities for each method. We then estimated the Pearson correlation coefficient between the suitability of the species in the communities and the probability obtained from each method. A good method should exhibit a strong correlation, reflecting its ability to characterize the suitability of each species in each community. We also examined the accuracy of each method (closeness to the 1:1 line) by estimating their mean absolute error (MAE):

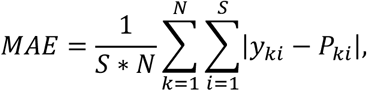

where *y*_*ki*_ is the real value of suitability for species *i* in site *k* and *P*_*ki*_ is the probability assigned by each method.

##### Test results

Our results revealed that the Hypergeometric method exhibited the most desirable characteristics. First, it showed the strongest correlation with suitability (which is ultimately the goal of a method for detecting dark diversity), followed by the Favorability method (Fig. 1). By contrast, the Beals index presented a substantially weaker correlation. However, Favorability exhibited a narrow range of predicted probabilities, with most values close to 0.5, despite suitability values were evenly spread across the entire 0-1 range. This resulted in Favorability being the least accurate method in our tests (MAE = 0.25). By contrast, the Hypergeometric method showed a much wider range of predicted probabilities, and more accurate estimations of suitability (MAE = 0.17; Fig. 1).

**Fig. 1.**
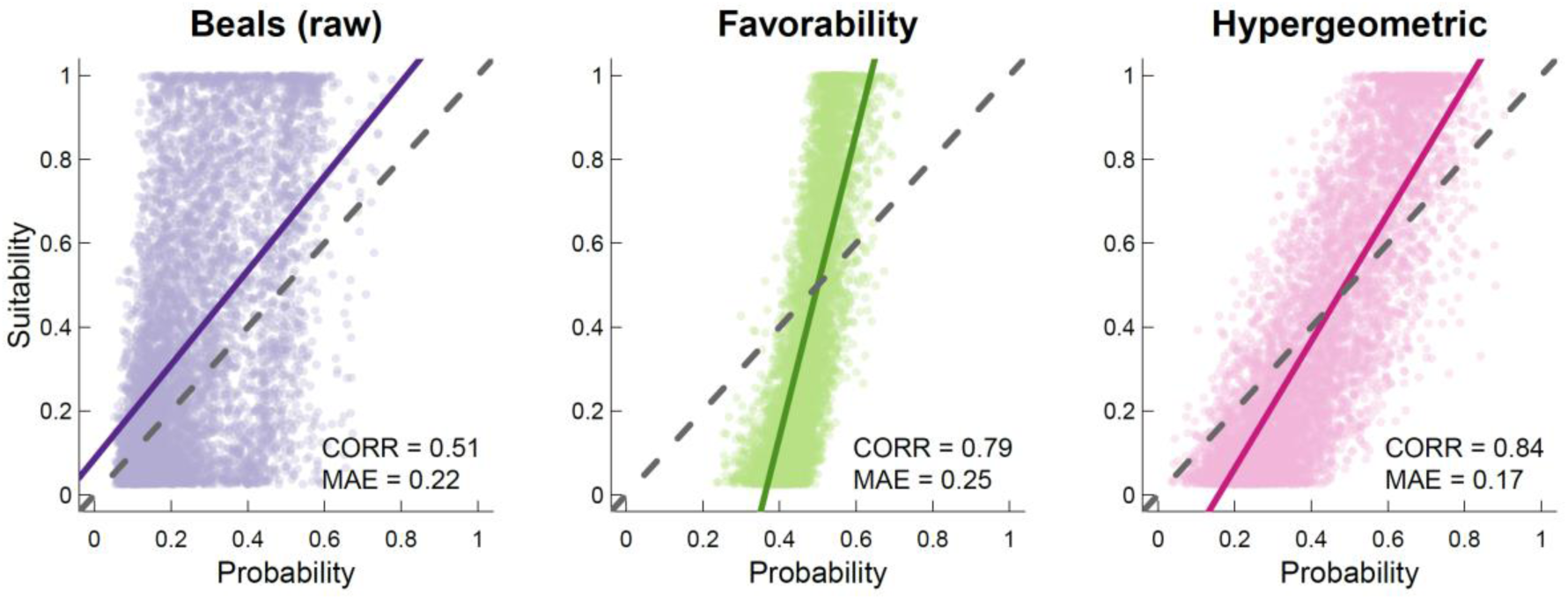
Relationship between the probabilities assigned by each method to the species absent from each community and their suitability in each community. Continuous coloured lines indicate the fit of a linear model between the two variables and the dashed line indicates a 1:1 relationship. Pearson correlation coefficient and mean absolute error (MAE; indicating closeness to the 1:1 line) are shown in each plot.

#### TEST 2: PREDICTIVE ABILITY AND RELATIONSHIP WITH DATASET SIZE

##### Test rationale

One potentially important aspect in comparing these methods is their sensitivity to the size of the dataset. Some methods may be more suitable for datasets containing many sites than those containing few sites. To examine this, we selected random subsets of varying size (from 5 to 95 communities in intervals of 5) of the communities after the last simulation step. From these reduced datasets, we estimated the correlation between the probability obtained from each method to absent species and their suitability in communities (as in Test 1). We repeated this procedure 100 times for each size, attaining 100 values of the correlation for each size and method. We then examined how the correlation improved as a function of the size of the dataset for each method. For this, for each subset (i.e. each sample size), we performed a linear mixed model using the method as a fixed effects explanatory variable and each random subset (100 repetitions) as a random effect. We then performed Tukey post-hoc tests to detect differences among methods.

##### Test results

Our results showed that the Hypergeometric method performed best for most sample sizes (Fig. 2). The Favorability method outperformed the Hypergeometric method only with a sample size of 5 communities, which is an unrealistically low value. The hypergeometric method’s performance increased more rapidly than that of the other methods with increasing number of communities and was the superior method for all sampling sizes greater than 15 communities. Beals’s predictive ability was inferior to Favorability’s for all sample sizes (Fig. 2).

**Fig. 2.**
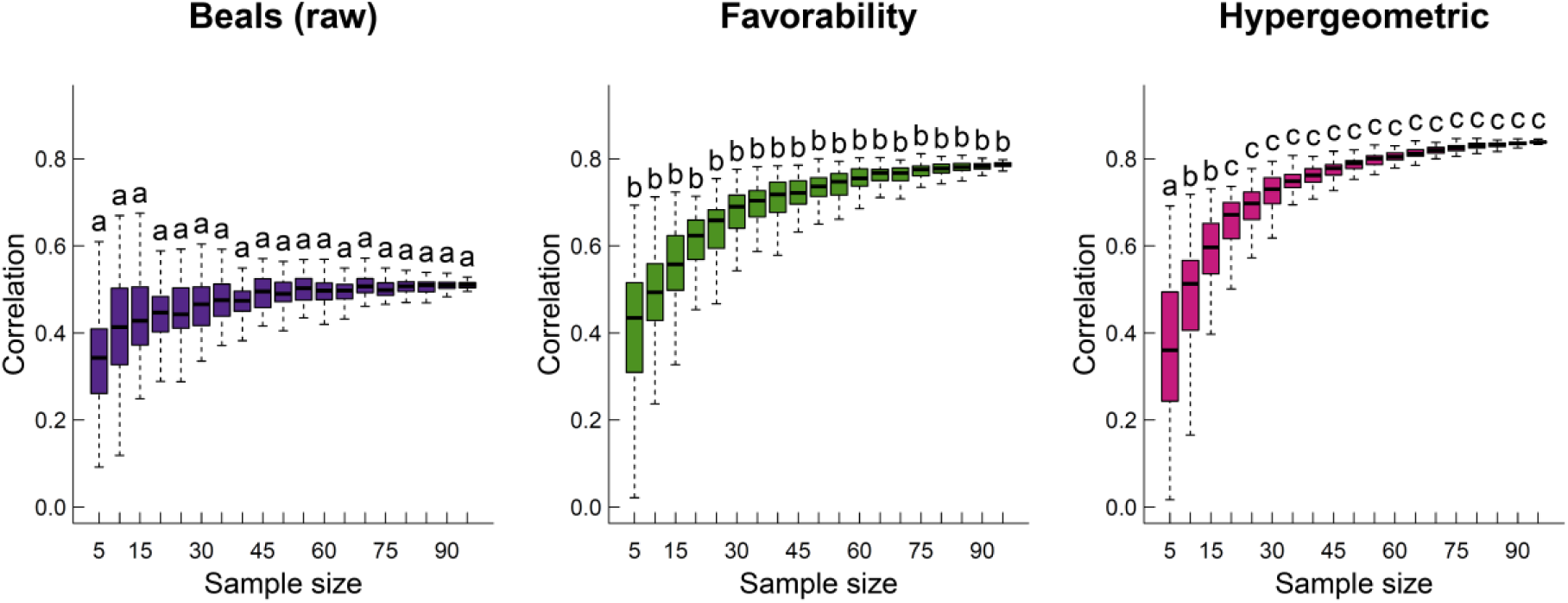
Predictive ability of the different methods as a function of sample size. Each plot shows how the correlation (Pearson) between the suitability of absent species in each plot and the probabilistic value given by each method varies as the number of plots increased (see main text for further explanations). Letters above each boxplot show differences in a Tukey post-hoc test (α=0.05) comparing methods within the same sample size, considering each random repetition as a random factor.

#### TEST 3: ESTIMATIONS OF DARK DIVERSITY SIZE

##### Test rationale

In some cases, it is interesting to characterize the size of dark diversity (i.e. its expected number of species). For this. the probabilities for all species in a given site can be added (Karger et al., 2016). This approach considers our level of certainty about species membership in dark diversity: species with low probabilities will count little towards the total dark diversity size, whereas species with high probabilities will contribute greatly. Using the data from the last simulation, we tested the relationship between the size of dark diversity predicted by each method and the sum of the suitability of the absent species in each community. As in Test 1, we also estimated MAE to assess the accuracy of each method.

##### Test results

The size of dark diversity based on the Beals index had a non-significant correlation with the size of dark diversity based on suitability (p = 0.485; Fig. 3). By contrast, both the Favorability and the Hypergeometric methods exhibited positive relationships between the predicted size of dark diversity and the size of dark diversity based on suitability, with similar predictive ability (p < 0.001 in both cases; Fig. 3). However, the sizes of dark diversity estimated with the hypergeometric method were much more similar to those based on suitability (59.3% reduction in MAE), whereas sizes based on suitability were always overestimated.

**Fig. 3.**
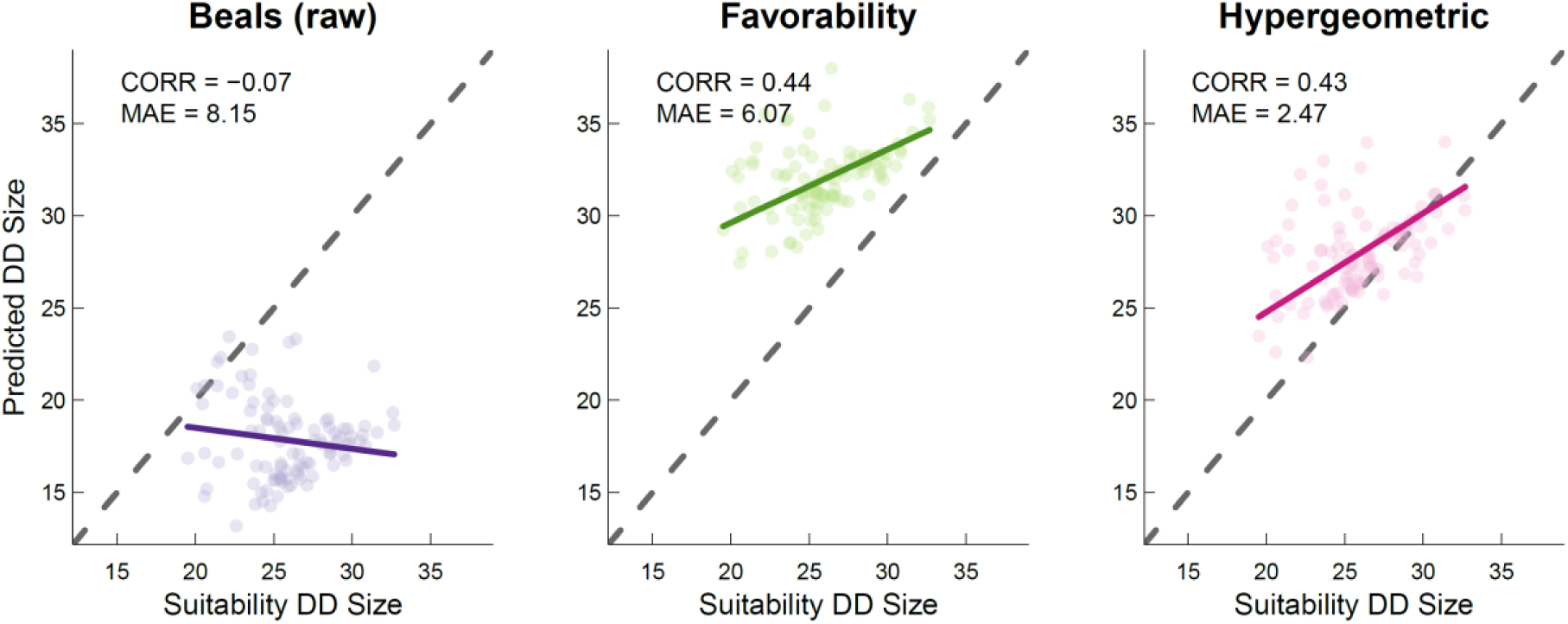
Relationship between the size of dark diversity predicted by each method and the true size of dark diversity according to the summed suitability of absent species in each community. Continuous coloured lines indicate the fit of a linear model between the two variables and the dashed line indicates a 1:1 relationship. Pearson correlation coefficient and mean absolute error (MAE; indicating closeness to the 1:1 line) are shown in each plot.

#### TEST 4: CORRELATION BETWEEN PREDICTIONS AND SPECIES REGIONAL FREQUENCY

##### Test rationale

Finally, we explored the effect of species regional frequency on the values that each method predicts. The predictions of a method that simply reflects species frequency will be biased (greater probabilities for more frequent species), and not satisfying the original definition of dark diversity, which does not depend on species regional frequency, but rather on their ecological requirements. To explore this, we estimated the correlation between the probability obtained for species not observed in the community and the frequency of species in the dataset (number of communities in which a species was found). Ideally, suitable methods to estimate probabilistic dark diversity should not show strong correlations between these two variables.

##### Test results

The predictions of the Beals index showed an extremely strong positive correlation with species regional frequency (Fig. 4). The other two indices also showed positive (but notably weaker) correlations with regional frequency. Favorability, in principle designed to mitigate this correlation, exhibited the weakest correlation, whereas the Hypergeometric method predictions were the least affected by the regional frequency of species (Fig. 4).

**Fig. 4.**
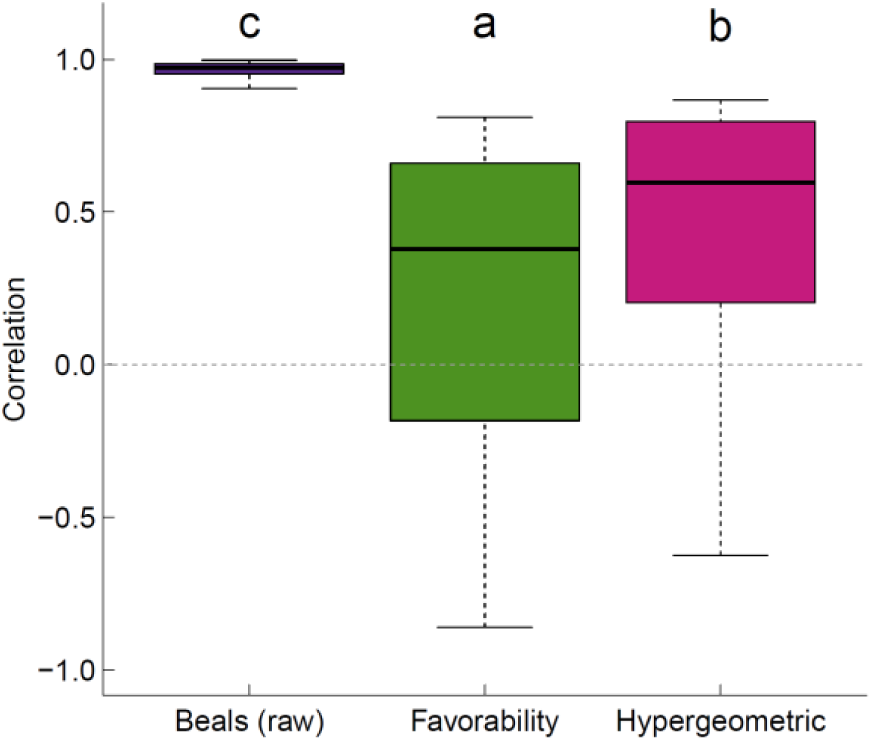
Correlation (Pearson) between the probabilities predicted by each method for the absent species from each site and the regional frequency of species in the simulated dataset. Letters above each boxplot show differences in a Tukey post-hoc test (α=0.05) comparing methods.

#### Real data example

We applied the three methods in two vegetation datasets with a nested hierarchical sampling design. The first dataset was a systematic sample of Swiss forests (“Swiss dataset”; Wohlgemuth, Moser, Brändli, Kull, & Schütz, 2008), with species recorded in 707 sites at two nested scales (30 m^2^ and 500 m^2^), with a total of 772 species. The second dataset contained coastal grassland vegetation from Scotland (“Scottish dataset”; Shaw, Hewett, & Pizzey, 1983), encompassing 3033 sites and 465 species. Species identities were also recorded at two nested scales (4 m^2^ and 200 m^2^). Following Lewis et al. (2015), we built species x species co-occurrence matrices in the smaller plots, and then estimated probabilistic dark diversity using the three different probabilistic co-occurrence based methods.

Similarly to Test 1 for the simulated dataset, we explored the probabilities obtained from each method for all species in all communities. We also compared the probabilities that each method assigned to species designated as “Absent” (species present in neither nested plots), as “Dark” (species absent from the small plot but present in the large plot), and “Observed” (species present in the small plot). Finally, as in Test 4 for the simulated dataset, we explored the correlation between the values predicted by each method and the frequency of the species in the region.

In these datasets, both the Hypergeometric and the Favorability methods predicted probabilities encompassing the whole 0-1 range, with average predictions being around 0.4 for both methods. By contrast, the raw Beals index predicted extremely low probabilities on average (Fig. 5). This behaviour reflects the effect of regional frequency in the Beals raw index; this effect was absent in the favorability correction, which should reflect deviations from the general frequency of species (thus being higher for sites where the conditions are better suited for the species; Real et al. 2017). The distribution of Favorability probabilities was bimodal, with one peak of probabilities equal to 0, much more marked in the Scottish dataset (30.3% of the predicted probabilities in the Scottish dataset and 12.7% in the Swiss dataset were exactly 0), and the second peak resembling a normal distribution centred around 0.5. This bimodality was caused by the great number of rare species occurring only in one or two sites, which generally are assigned a 0 probability in the Beals index method, and which is maintained in the Favorability method. By contrast, the Hypergeometric method assigned to these species in most cases a probability slightly less than 0.5. The Hypergeometric method produces probabilities near 0.5 in two situations: from a genuine lack of association among species, or from a lack of information due to the species having low frequency (or theoretically high frequency). The latter restricts the number of ways in which species can co-occur (two species with only one appearance each in a dataset can co-occur in one site), and hence departures from random co-occurrences can never be large. Values close to 0.5 effectively express a lack of information on the ecological requirements of rare species, which can be considered an advantage of the Hypergeometric method. All methods worked similarly well in assigning ordered probabilities to species according to their status, with each method assigning the lowest probabilities to absent species and the highest probabilities to present species.

**Fig. 5.**
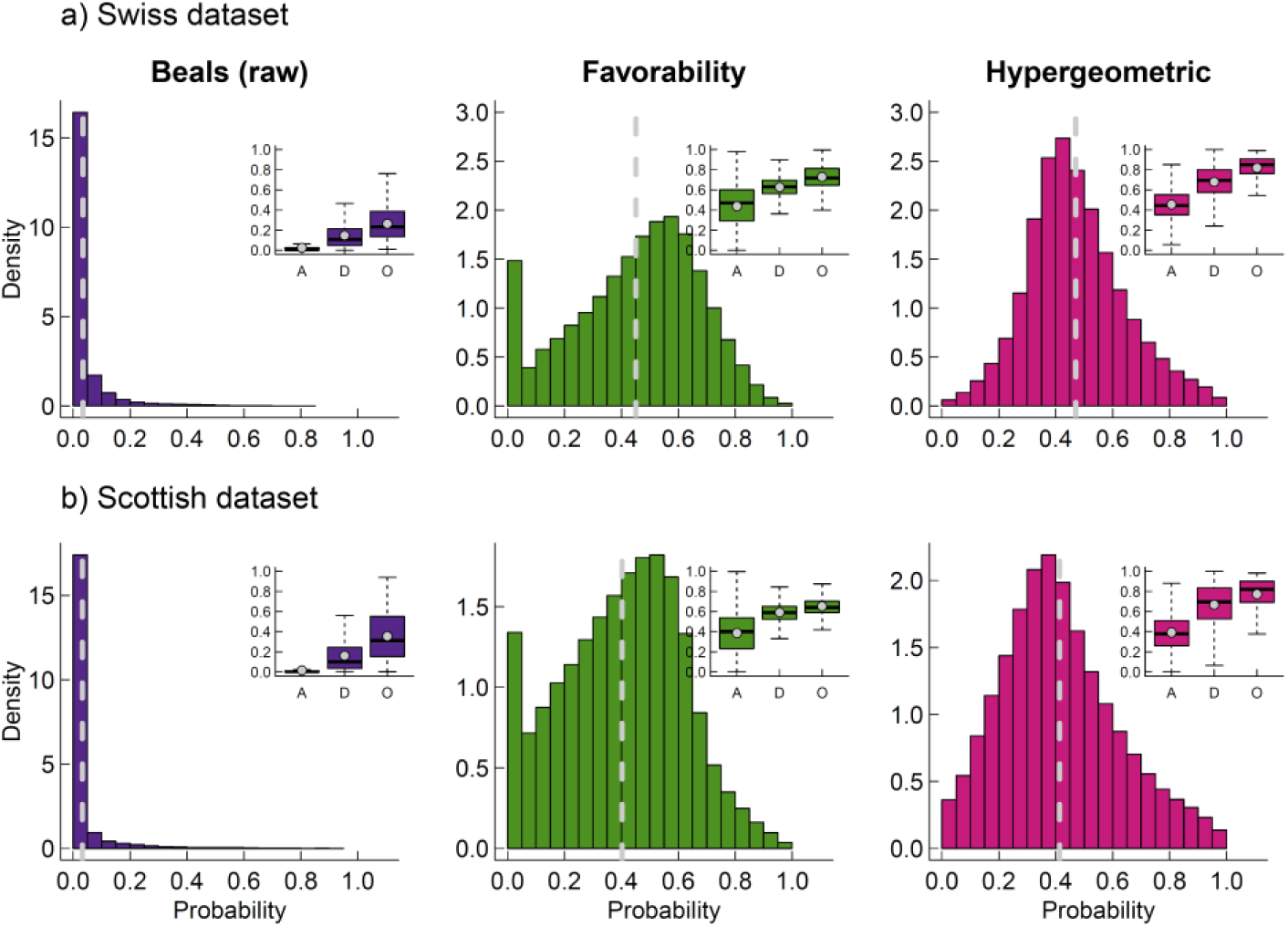
Distribution of the probabilities obtained from each method, considering all species in all the sites of each dataset. The grey dashed line indicate the average probability of each method in each dataset. The subplots show the different probabilities obtained from each method in each dataset to species categorized as “absent” (A; species not found in the considered site at any scale), “dark” (D; species found in the large plot, but not in the small one) and “observed” (O; species found in the small plot).

Similarly to the results of Test 4, the correlation between predictions of the Beals index and the regional frequency of species were extremely high (Fig. 6). In contrast with the simulated dataset, the Hypergeometric method—not Favorability—exhibited the weakest correlation, particularly in the Scottish dataset (Fig. 6), probably due to the aforementioned effect of rare species on Favorability predictions. Examination of the relationship between the frequency of the species and its average probability for the Hypergeometric method revealed probabilities close to 0.5 for the least frequent species, with the predictions becoming more variable as frequency increased.

**Fig. 6.**
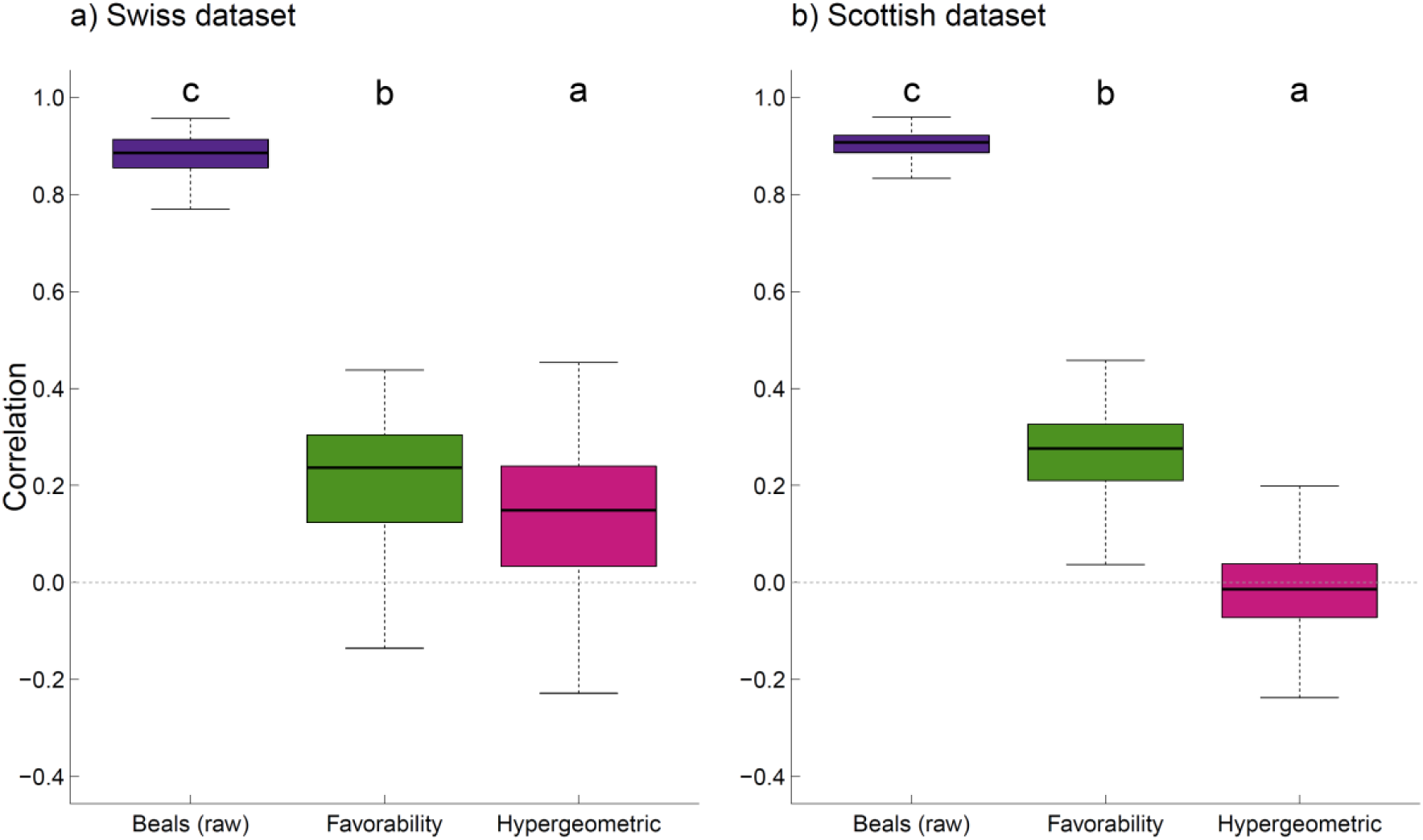
Correlation (Pearson) between the probabilities predicted by each method for the absent species from each site and regional frequency of species in the real datasets. Letters above each boxplot show differences in a Tukey post-hoc test (α=0.05) comparing methods.

## Discussion

With this study we aimed to advance the development of probabilistic methods to estimate dark diversity (the absent part of the site-specific species pool) using species co-occurrences. By linking local and regional scales, dark diversity can help us to understand better biodiversity and its dynamics (Pärtel, Bennett, & Zobel, 2016). However, unlike observed diversity, dark diversity is not directly measurable, and depends on algorithmic estimation. Here, we presented a fully probabilistic method to estimate dark diversity using the co-occurrence matrix of species based on the hypergeometric distribution (Griffith et al., 2016). We compared its performance with other two extant methods based on species co-occurrences (Beals and Favorability) using simulations that include information on the ecological affinities of species within communities (suitability). By considering several criteria (distribution of predicted probabilities, predictive ability, and estimations of dark diversity size) we found that, although Favorability was generally superior than Beals, the Hypergeometric method performed better than the two other probabilistic methods. Further, we compared the results obtained from each method in two real datasets, showing that the positive features of the Hypergeometric method are also apparent in real-world applications.

The fact that dark diversity cannot be observed directly has two important implications. First, by acknowledging this lack of determinism, probabilistic approaches are particularly attractive alternatives to estimate dark diversity (Real et al., 2017). Despite insistences that dark diversity should be estimated probabilistically have accompanied the concept since its inception (Mokany & Paini, 2011), only recently have such approaches been adopted (Brown et al., 2019; de Bello et al., 2016; Karger et al., 2016; Lessard et al., 2016). Second, measuring dark diversity poses a methodological challenge, since there are no appropriate benchmarks to compare methods. Previous tests of dark diversity estimation methods have used nested datasets or repeated sampling in order to “observe” dark diversity (Brown et al., 2019; de Bello et al., 2016; Karger et al., 2016; Lewis et al., 2016). These studies have frequently found that Beals is the most suitable method. Although some of these studies acknowledge the imperfection of these tests because only an unknown portion of the true dark diversity is observed (Brown et al., 2019), the observed part of dark diversity is non-random. This is because species that are found in the observed portion of the dark diversity of a site are likely to be not only ecologically suitable, but also to have a greater frequency in the region (De Cáceres & Legendre, 2008; Real et al., 2017). Although many highly suitable species might be absent due to dispersal limitation (Riibak et al., 2015; Zobel, 2016), species with high frequency in the region may have also a high availability of propagules and can be present in less suitable sites due to source-sink dynamics (Pulliam, 2000). As a consequence, using nested or resampled datasets to calibrate and compare methods to estimate dark diversity can lead to biased results favouring indices—such as the Beals index—that predict greater probabilities for the most frequent species. Although the Beals index is a good predictor of the probability of occurrence of the target species (De Cáceres & Legendre, 2008), it is not necessarily a good predictor of their suitability in a given site. At this point it is important to consider that—despite the different definitions attributed to “species pool” (Zobel, 2016)—dark diversity refers to those species that are absent from a site despite suitable ecological conditions (Pärtel et al., 2011). According to this criteria, our simulations revealed that Beals was clearly outperformed by the two other methods in terms of its ability to estimate suitability and dark diversity size. Consequently, while we agree that the raw Beals index can be useful for predicting which species will be observed as we increase sampling effort (either in space or in time; Karger et al., 2016), this is largely because it serves as a very good proxy of species general frequency, and more frequent species are found more often. However, we recommend that future studies estimating species adscription to dark diversity should focus on the ecological affinities of species, rather than on predicting occurrences in space or time.

Favorability and Hypergeometric methods are less affected by species frequency, and better indicators of the site suitability. Favorability, based on a correction of Beals to remove the effect of species frequency (Real et al., 2017), predicted species suitability and dark diversity size in our simulations better than Beals. However, the method was not completely free from the effect of species frequency in the region, since it assigned 0 probability to species for which there was no information (i.e. none of the species recorded in the site had co-occurred with the target species), which tended to be extremely rare species. Such extreme predictions for species with little information is not what one would expect if probabilities of adscription to dark diversity reflect the suitability of species in a site. This is not an issue of the Favorability transformation itself, but is inherited from the fact that the Beals index can result in probabilities of exactly 0. By contrast, the Hypergeometric method assigned probabilities close to 0.5 in these rare species, thereby expressing better the lack of available information: whether to include very infrequent species in dark diversity is akin to coin flipping, whereas more confident predictions can be made for common species. In fact, the Hypergeometric method is most reliable for pairs of species with intermediate incidence (Lavender, Schamp, Arnott, & Rusak, 2019). In any case, the Hypergeometric method outperformed Favorability in all considered aspects of our simulations. Although all methods proved capable of discriminating between observed and non-observed species, the distribution of probabilities of the Hypergeometric method exhibited the most appropriate shape, encompassing the whole range of available probability. Moreover, it was the best method for predicting the ecological affinity of species for all reasonably sized datasets. It was also the best calibrated method, returning unbiased predictions of both suitability and dark diversity size. In addition, it exhibited positive features in real datasets, including the aforementioned lack of extreme predictions, a good ability to resolve between absent and present species, and a reasonable relationship between predicted probabilities and species frequency. As a consequence, we conclude that the Hypergeometric method is currently the most promising method to estimate probabilistic dark diversity and species pool composition based on co-occurrences.

## Conclusions

Methods based on species co-occurrence patterns have proven to be a powerful tool to estimate probabilistic dark diversity. They integrate information on abiotic and abiotic conditions, which makes them good at characterizing the realized niches of species (Lewis et al., 2016). Most importantly, information on species co-occurrences is increasingly available in a wide range of environments and regions, which should allow us to improve estimation of species pairwise associations. An important aspect to consider is that correct characterizations of dark diversity based on species co-occurrences require reliable and complete sampling of the species that are present. This can be challenging for sites containing many elusive or inconspicuous species (Boussarie et al., 2018). On the other hand, estimations of probabilistic methods might help to improve assessment of observed diversity by indicating apparently absent species with a high probability of having eluded detection. Among the methods considered here, existing evidence suggests that the Hypergeometric method is the most suitable to detect pairwise associations among species (Lavender et al., 2019). However, species do not occur in pairs, but form diverse interacting networks, so that restricting our analyses to pairwise co-occurrences is likely neglecting substantial amounts of ecological information. Future methods to estimate probabilistic dark diversity would benefit greatly from co-occurrence based methods that look beyond associations between pairs of species. Considering biodiversity from a probabilistic point of view is a meaningful way to acknowledge the uncertainty associated with ecological information. The development of probabilistic dark diversity joins similar advances made in functional diversity (Carmona, de Bello, Mason, & Leps, 2016). Future integration of probabilistic species pools and functional diversity will advance our understanding of assembly processes and conservation status of ecological systems at multiple spatial and temporal scales. In order to help ecologists implement all the methods shown here, we have developed the ‘DarkDiv’ R package (Carmona, 2019; freely available in https://CRAN.R-project.org/package=DarkDiv).

